# Wax bloom dynamics on *Sorghum bicolor* under different environmental stresses reveal signaling modules associated with wax production

**DOI:** 10.1101/2024.10.10.617702

**Authors:** Madison Larson, Marshall Hampton, Lucas Busta

## Abstract

Epicuticular wax blooms are associated with improved drought resistance in many species, including *Sorghum bicolor*. While the role of wax in drought resistance is well known, we report new insights into how light and drought dynamically influence wax production. We investigated how wax quantity and composition are modulated over time and in response to different environmental stressors, as well as the molecular and genetic mechanisms involved in such. Gas chromatography-mass spectrometry and photographic results showed that sorghum leaf sheath wax load and composition were altered in mature plants grown under drought and simulated shade, though this phenomenon appears to vary by sorghum cultivar. We combined an *in vitro* wax induction protocol with GC-MS and RNA-seq measurements to identify a draft signaling pathway for wax bloom induction in sorghum. We also explored the potential of spectrophotometry to aid in monitoring wax bloom dynamics. Spectrophotometric analysis showed primary differences in reflectance between bloom-rich and bloomless tissue surfaces in the 230-500nm range of the spectrum, corresponding to the blue color channel of photographic data. Our smartphone-based system detected significant differences in wax production between control and shade treatment groups, demonstrating its potential for candidate screening. Overall, our data suggest that wax extrusion can be rapidly modulated in response to light, occurring within days compared to the months required for the changes observed under greenhouse drought/simulated shade conditions. These results highlight the dynamic nature of wax modulation in response to varying environmental stimuli, especially light and water availability.

**Significance Statement:** Agricultural crops require significant freshwater for irrigation, making food security vulnerable to drought. Epicuticular wax blooms are associated with drought tolerance in many plants, including *Sorghum bicolor*. We investigated how environmental factors like light and drought influence wax production in sorghum. Wax production, composition, and gene expression were compared between sorghum exposed to different environmental stressors, reavealing dynamic modulation of wax production in response to environmental stress as well as signaling genes potentially involved in regulating wax production. These findings broaden our understanding of wax-related drought tolerance mechanisms, providing a foundation for future efforts to enginner crops with improved climate resilience.

## 1. Introduction

Understanding the chemical and biological mechanisms underlying crop resilience and productivity is crucial for addressing the challenges faced by modern agriculture. Global food production is often limited by the availability of water, making irrigation a crucial resource and high-intensity droughts a major challenge. Substantial freshwater use is associated with irrigation of key crops like maize, legumes, and alfalfa (1). The reliance of agricultural production on water availability underscores the potential for food security to be jeopardized by drought, such as that experienced by much of the Midwestern USA during the summer of 2021 (2). The mechanisms underlying drought resistance have been the focus of extensive research, yet our understanding remains incomplete (3–21). Failing to expand our understanding of drought tolerance mechanisms risks leaving crops vulnerable to the escalating challenges posed by climate change and water scarcity.

While maize and sorghum are both C4 cereal grasses that share similar vegetative phase morphology, they exhibit distinct mechanisms for combating drought stress. One drought tolerance mechanism adapted by sorghum and other drought tolerant species from a variety of genera is the production of thick epicuticular wax coatings (wax blooms) (22). These waxes are associated with improved water use efficiency (23–27). In several species, including sorghum, changes in wax bloom production have been observed following exposure to environmental stressors like drought and light intensity (25, 26, 28–45). Given that wax blooms are associated with drought tolerance, and maize (a more economically and nutritionally relevant crop) lacks a thick epicuticular wax layer, understanding the molecular basis for their occurrence in sorghum is valuable. The mechanisms of wax biosynthesis have been studied extensively (4, 46–53), however, the signaling pathways regulating wax bloom production in response to environmental stimuli like water or light availability are not well understood.

The objective of this study was to improve our understanding of how wax synthesis and transport (quantity and/or composition) is modulated over time and in response to different environmental stressors, as well as the molecular and genetic mechanisms underlying this modulation. To achieve this goal we grew sorghum under varied environmental conditions while monitoring and quantifying wax blooms via photographic and chemical measurements, then analyzed the corresponding data to determine if wax bloom production was affected by any experimental stimuli. We also performed *in vitro* wax induction experiments from which we gathered wax load data to validate the inducibility of wax production. We also collected RNA-seq data from wax induction experiments and extensively integrated it with existing literature to form a hypothesis for a light signaling pathway mediating the induction of wax production. This work represents a step towards expanding our fundamental knowledge of a drought tolerance mechanism that could help reduce the impacts of climate change on agriculture.

## 2. Materials and Methods

### A. Plant material and growth conditions, and in vitro wax induction

Germination: Sorghum bicolor, var. BTX623 seeds were germinated in three inch pots filled with a soil mixture containing one part Promix ®BX general propagation mix and one part perlite under LED grow lights (Feit Electric GLP36FS) with a light intensity of approximately 340 lux (measured using IOS application “Light Meter”). After germination, plants were transplanted into twelve inch pots, with two plants per pot. Following transplantation, all plants were watered once every seven days using a fertilizer solution containing one teaspoon of Jack’s Classic All Purpose Fertilizer per gallon of water until experimental treatments began. All plants were kept under identical light conditions as well until experimental conditions were initiated. Experimental conditions were initiated once all plants had at least 8 fully emerged leaf sheaths. Light conditions were as follows: Plant Room (germination): 340 lux, Greenhouse (control), Greenhouse (low shade): 900 lux, Greenhouse (high shade): 650 lux. Control, light and water treatment groups included four, two and six plants respectively. For light manipulation treatments control plants were grown under full sun on the greenhouse benchtop, and treatment groups were grown inside structures covered by one and two layers of shade cloth, respectively. Water treatments were structured such that control plants received weekly waterings, while treatment groups received water every nine days and eleven days, respectively.

Induction of epicuticular waxes on sorghum leaf sheath tissue was performed according Jenks et al., 1994 using a light intensity of approximately 340 lux (54).

### B. Analysis of epicuticular wax blooms with gas chromatography-mass spectrometry and spectrophotometery

Sorghum leaf sheath segments were first photographed, then added to scintillation vials and surface waxes were extracted by rinsing twice with HPLC grade CHCl3 (MilliporeSigma) for approximately 30 seconds. The resulting solutions were transferred to GC vials, 25 *μ*L 0.46mg/mL tetracosane standard (C24H50, Alfa Aesar) were added, and excess reagents were evaporated overnight. Samples were derivatized in 50 *μ*L of 1:1 N,O-Bis(trimethylsilyl)trifluoroacetamide (BSTFA; Restek Corporation) and pyridine (Acros Organics) and incubated (70°C for 45 min.). Samples were analyzed using a 7890B Network GC (Agilent) and 7693A Autosampler (Agilent) equipped with a split/splitless injector and an HP-5 capillary column (Agilent, polydimethylsiloxane, length 30 m x 0.320 mm, 0.10 *μ*m film thickness). 2 *μ*L of sample was injected into a constant flow of He maintained at 1.4mL/min. The GC oven temperature was held at 50°C for 2 min, then increased by 40°C/min to 200°C, held at 200°C for 2 min, then increased by 3°C/min to 320°C and held at 320°C for 30 min. The total run time was just over 77 minutes and a solvent delay of eight minutes was used. Wax components were detected using an Agilent 5977B (GC/MSD) mass selective detector (EI 70eV; m/z 40-800, 1 scans/s).

Reflectance measurements were collected for sorghum leaf sheath tissue both with and without visible wax blooms using a double beam spectrophotometer (Cary 5000) with an external diffuse reflectance accessory. Waxless tissue was the same tissue used for wax-covered measurements with the wax bloom physically removed using a paper towel.

### C. Differential Expression Analysis

Sorghum RNA was sequenced using the Illumina platform at the University of Minnesota Genomics Center (250bp read legnth, 20M paired reads per sample). Transcript counts were quantified using Kallisto and differential expression analysis was performed using the R package DESeq2. Maize gene expression data were obtained from the NCBI SRA database (SRA accession numbers: SRR5985066, SRR5985076, SRR5985077, SRR5985051, SRR5985056, SRR5985070).

### D. Photographic data collection and analysis

Photographic measurements were performed on the tenth leaf sheath of sorghum plants beginning five days after emergence of the leaf sheath tip. Each photograph included a QR code containing the experimental information corresponding to each experimental plant, as well as a colorchecker (Calibrite) device to enable categorization and normalization of photographic data using our analysis application. Photographs were stored in google drive folders labeled by collection date for future additional analysis. Code for the analysis application and all other analyses can be found at https://github.com/thebustalab/wax_bloom_dynamics/.

## 3. Results

The objective of this study was to improve our understanding of: (i) how wax extrusion (quantity and composition) is modulated over time and in response to different environmental stressors, and (ii) the molecular and genetic mechanisms regulating wax bloom extrusion. To meet these objectives, we first used GC-MS to monitor wax quantity and composition on sorghum plants grown under drought or shade conditions (section 3A). We then investigated gene expression during the light induction of sorghum leaf sheath wax blooms using a previously published wax induction protocol, RNA-sequencing, and differential expression analysis (section 3B) (54). Finally, we began laying the groundwork for high throughput screening of candidate wax signaling genes using comparative genomics and a phenotyping apparatus we developed (section 3C).

### A. Sorghum leaf sheath wax extrusion and wax composition under drought and simulated shade stress

To investigate the dynamics of wax bloom extrusion under the influence of different environmental stressors, gas chromatography-mass spectrometry was used to quantify wax abundance on sorghum leaf sheaths grown under control conditions, as well as conditions of high simulated shade (650 lux), low simulated shade (900 lux), low (intensity) drought (watered every 9 days) and high (intensity) drought (watered every 11 days; Fig. 1A). Average wax coverage for plants grown under control conditions was 38 ± 9 *μg cm*^−2^, and average wax coverages on low intensity drought (34 ± 11 *μg cm*^−2^) and shade (27 ± 12 *μg cm*^−2^) treatment groups were not significantly different from the control group (Tukey test, p-values ≤ 0.05). However, average wax coverages differed significantly from the control group under high intensity drought (higher wax coverage, 64 ± 18 *μg cm*^−2^) and high intensity simulated shade (lower wax coverage, 17 ± 5 *μg cm*^−2^).

**Fig. 1.**
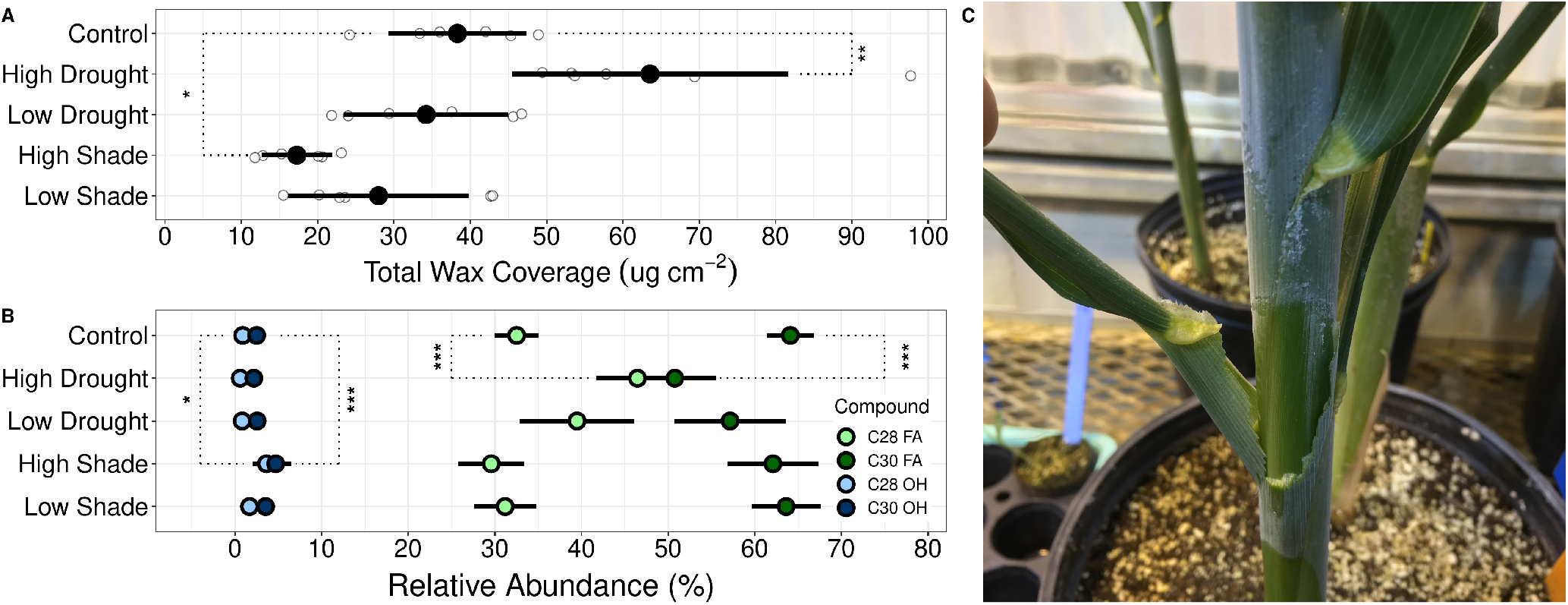
Sorghum leaf sheath wax extrusion under drought and shade stress. **A** Chemical response of sorghum wax blooms to light and water availability (Tukey test, * p-value ≤ 0.05, ** p-value ≤ 0.01). White points represent individual measurements. Black points and error bars represent average and standard deviation of n = 6 replicates, respectively. **B** Wax composition for sorghum grown under limited light or water conditions (* p-value ≤ 0.05, *** p-value ≤ 0.001). Colored points and error bars represent average and standard deviation of n = 6 replicates, respectively. **C** Sorghum plant with leaf sheath partially removed to reveal bloomless subtending leaf sheath tissue. **D** Shade cloth structures used for simulated shade treatments.

In addition to wax coverage, gas chromatography-mass spectrometry was also used to quantify and compare wax composition between sorghum plants grown under the control and drought/shade conditons. Similar to the trends in our wax load data (Fig. 1A), there were significant differences in wax composition between the plants in the control group and plants in the high intensity treatment groups, but not low intensity treatment groups (Tukey test, p-values ≤ 0.05). High intensity drought treatments led to significant increases in C28 fatty acid abundance and significant decreases in C30 fatty acid abundance compared to control (Fig. 1B). The average abundances of C28 and C30 fatty acids for the control group were (32 ± 3%) and (64 ± 3%), respectively, while the average abundances of C28 and C30 fatty acids for the high intensity drought treatment group were (46 ± 5%) and (51 ± 5%), respectively. High intensity shade treatments led to significant increases in both C28 and C30 alcohols compared to control, where the average abundances of C28 and C30 alcohols for the control group were (1 ± 1%) and (2 ± 1%), respectively, while the average abundances of C28 and C30 alcohols for the high intensity shade treatment group were (4 ± 2%) and (5 ± 2%), respectively. Thus, high intensity environmental treatments both affected wax coverage and composition while low intensity treatments did not, with the most substantial effect being that of the high intensity drought treatment, which increased wax coverage on leaf sheaths by more than 50%.

### B. Light-based induction of sorghum wax blooms and associated changes in gene expression

While monitoring sorghum in the greenhouse during the drought and simulated shade experiments described above, we noticed that tissue which was covered by, or had recently emerged from beneath, overlying tissue displayed little to no visible wax accumulation, while tissue which had been uncovered for longer durations displayed substantial visible wax accumulation (Fig. 1C). We hypothesized that light could be a key stimulus to which overlying and subtending tissues respond differently to yield the differences in wax accumulation we observed. This hypothesis prompted us to use light exposure as a means to investigate the molecular mechanisms underyling the modulation of sorghum wax bloom production observed under high intensity drought and light stress. To test our hypothesis, we implemented a published system for light-mediated wax bloom induction that relies on exposing excised sorghum leaf sheath tissue segments to light (54). Sorghum leaf sheath segments were harvested from beneath 3-4 layers of superseding tissue, then some segments were exposed to light, while others were kept wrapped in foil (Fig. 2A). After 72 hours, the light-exposed tissue had accumulated a visible wax coating that, by GC-MS analysis, was composed of, on average, 3.5x more wax than was present on the glossy surfaces of the plants kept in the dark (Fig. 2B). Average wax accumulation for light-maintained samples was 14 ± 6 *μg cm*^−2^ and average wax accumulation for control samples was 4 ± 2 *μg cm*^−2^. Thus, on top of providing a highly controllable source of tissue for studying wax bloom induction, this experiment further confirmed the role of light in sorghum epicuticular wax bloom induction, as we initially found in our greenhouse study (Fig. 1A).

**Fig. 2.**
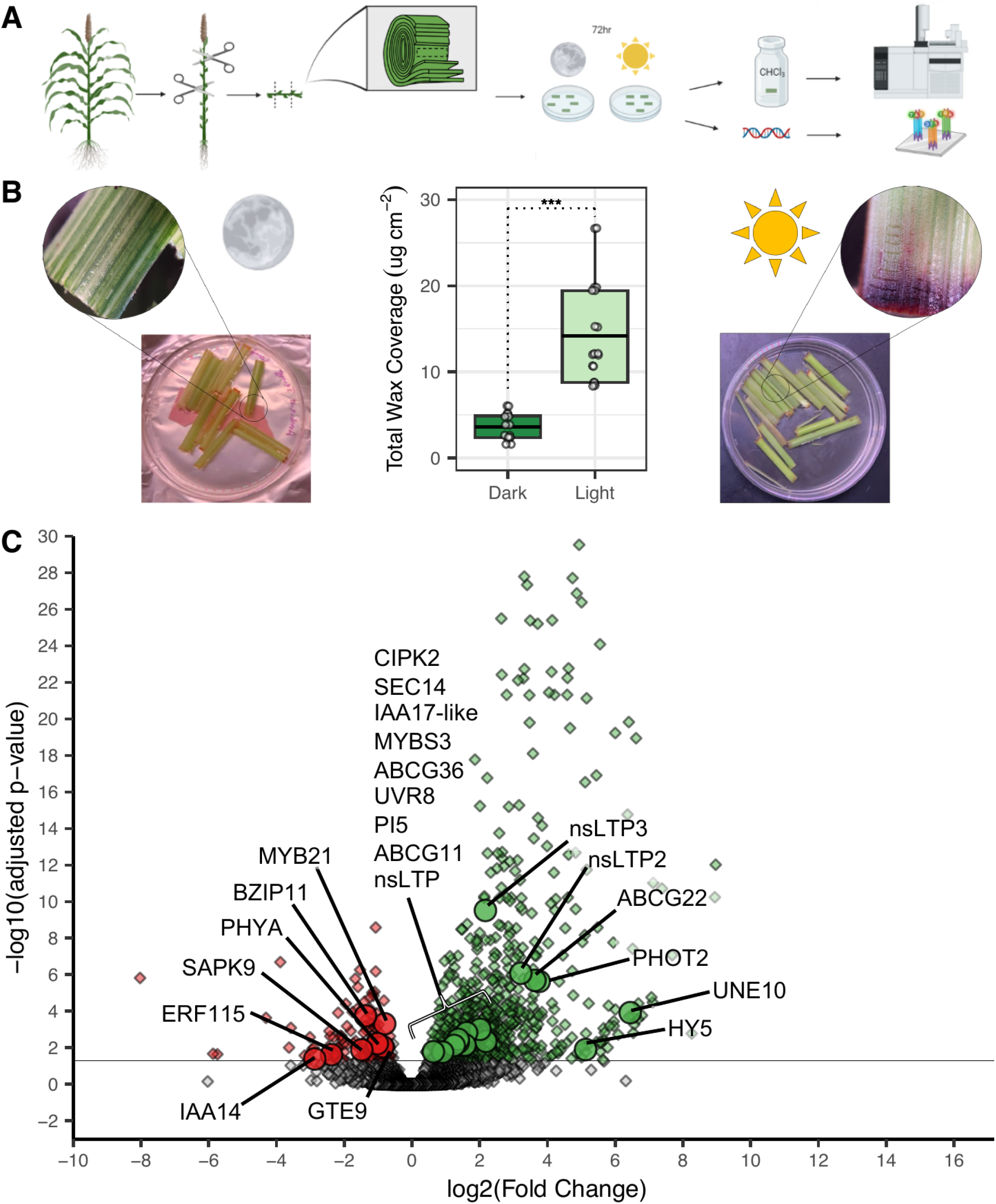
Impact of exposure to light on gene expression and wax chemistry of excised sorghum leaf sheath tissue segments. **A** Schematic of the light induction of wax bloom experimental workflow based on the protocol described by Jenks et al., 1994. Created with Biorender.com. **B** Gas chromatography-mass spectrometry measurements of wax coverage on sorghum tissue segments maintained in light versus complete darkness (T-test p-value ≤ 0.05). The line in the middle of each box of the box plot represents median total wax amount on respective tissues. **C** Volcano plot representing −log10(Padj) versus log2(Fold Change) for a differential expression analysis performed on transcriptomic data from sorghum tissue maintained in light versus complete darkness (FDR = 0.05). Red points represent genes with log2fold change ≤ (0.65), green points represent transcripts with log2fold change greater than or equal to (0.65).

So far, our results had shown that light as well as drought influence sorghum leaf sheath wax chemistry, and that exposure to light can induce the production of epicuticular wax blooms in excised sorghum tissue. A previous study of maize cuticle development also identified light as a key influencer of maize leaf wax chemistry (55, 56). Interestingly, while light stimulus causes both maize and sorghum to alter their leaf wax chemistry, it causes sorghum to produce massive epicuticular wax blooms that, even under various light treatments, do not occur on maize plants. We wanted to understand the genetic basis for this difference in chemical phenotype, and so we attempted to identify genes that were upregulated during sorghum wax production but whose orthologs were not upregulated during maize’s response to light. A previous study performed a detailed investigation of gene expression in maize leaf tissue as it is exposed to light, providing a reference against which to compare transcriptional responses in sorghum under light exposure (57).

To study the molecular mechanisms underlying the observed differences in wax bloom extrustion exhibited by sorghum tissue maintained under different light conditions, we examined gene expression in light- and dark-maintained excised sorghum leaf tissue as described in the light induction experiments described above (as shown in Fig. 2A). RNA from light- and dark-maintained sorghum tissue was extracted and sequenced, then sequencing reads were then mapped to a sorghum reference genome (V3.1) and transcript abundances were quantified. A differential expression analysis was used to identify genes that were up or downregulated in response to light, as wax production was being induced. We found that 825 genes were differentially expressed in response to light, with 693 genes being upregulated, and 132 being downregulated. We considered these genes as candidates that may be involved in light-mediated induction of wax blooms. Contrary to what was expected, at 72 hours post light exposure, we did not detect the upregulation of classic wax biosynthesis genes, including fatty acyl-CoA elongase complex genes like KCS, KCR, HCD, ECR, and CER2 as well as fatty acyl-CoA modification genes like CER1, CER3, CER4, and WSD genes, though we did observe the upregulation of some potentially biosynthesis-related esterase-like enzymes. Instead, our differentially expressed genes included photoreceptors (phytochrome A (phyA), cryptochrome DASH (CRY-DASH), phototropin 2 (PHOT2), UVR8), light signaling genes (HY5, phytochrome interacting factors), transcription factors (MYB, bHLH, bZIP), hormone related genes (IAA14, CBL-interacting kinase, GTE9, SAPK9, bZIP11, ERF115), and wax transport-related genes (acyl-coA binding proteins, lipid transfer proteins, ATP-binding cassette transporter subfamily G, SEC14). These candidate genes can serve as a starting point for future researchers aiming to expand our understanding of the regulatory mechanisms underlying wax blooms in sorghum, and we discuss further details of potential wax induction signaling methods below.

### C. Wavelength-specific reflectance of sorghum wax blooms and their tracking with smartphone-based phenotyping

Our investigation of changes in gene expression in response to wax bloom-inducing conditions revealed a substantial number of differentially expressed genes potentially involved in signaling the extrusion of wax bloom. The high number of candidates identified in our expression data combined with the candidates already described in literature suggest that high-throughput screening methods for monitoring sorghum wax bloom extrusion dynamics are needed if we hope to use diverse sorghum lines, or mutant lines to study the regulatory landscape underlying the light-inducibility of sorghum wax blooms. Since wax-covered and wax-less sorghum leaf sheath tissues can be detected with the naked eye (Fig. 3A), we hypothesized that it might be possible to monitor dynamic changes in wax coverage using a smartphone camera. To test this hypothesis, we started with analyses of sorghum photographs and reflectance data collected with a spectrophotometer to determine the which spectral regions might be most useful in monitoring wax load. From these analyses, we determined that the greatest difference in reflectance between wax-covered and wax-less tissue occurs from ~230-500nm suggesting that light in the UV/blue region of the spectrum is reflected more strongly by epicuticular waxes than other wavelengths in the spectrum (Fig. 3B). We also collected extensive photographic data from 20 sorghum plants grown under control conditions as well as high and low drought and simulated shade conditions by photographing corresponding internodal sections of each plant every other day before, during, and after wax extrusion. We used R code to systematically analyze the colorimetric data in the resulting 1360 photographs, resulting in a red, green, and blue color channel values for every pixel of every photograph, a total of 12,155,286 data points. We conducted a principal components analysis of the photographic data from controls plants and it showed that showed that on young, wax-less sorghum plants the changes in our photographic data were primary associated with the red and green color channels (Fig. 3C). However, as the plants matured and wax started accumulating on their leaf sheaths, the blue color channel of the RGB photographic data begins to change substantially more than the green and red channels (Fig. 3C). These results suggested that changes in wax coverage are more easily detected in the blue color channel of photographic data than the red and green channels. Accordingly, we propose that the blue color channel of photographic data could be used for monitoring wax accumulation with photographs. Using this application to analyze the photographic data from greenhouse experiments, we were able to detect significant differences in wax accumulation between the control and both low and high intensity shade treatment groups but were unable to detect significant differences between control and drought treatment groups (Fig. 3D).

**Fig. 3.**
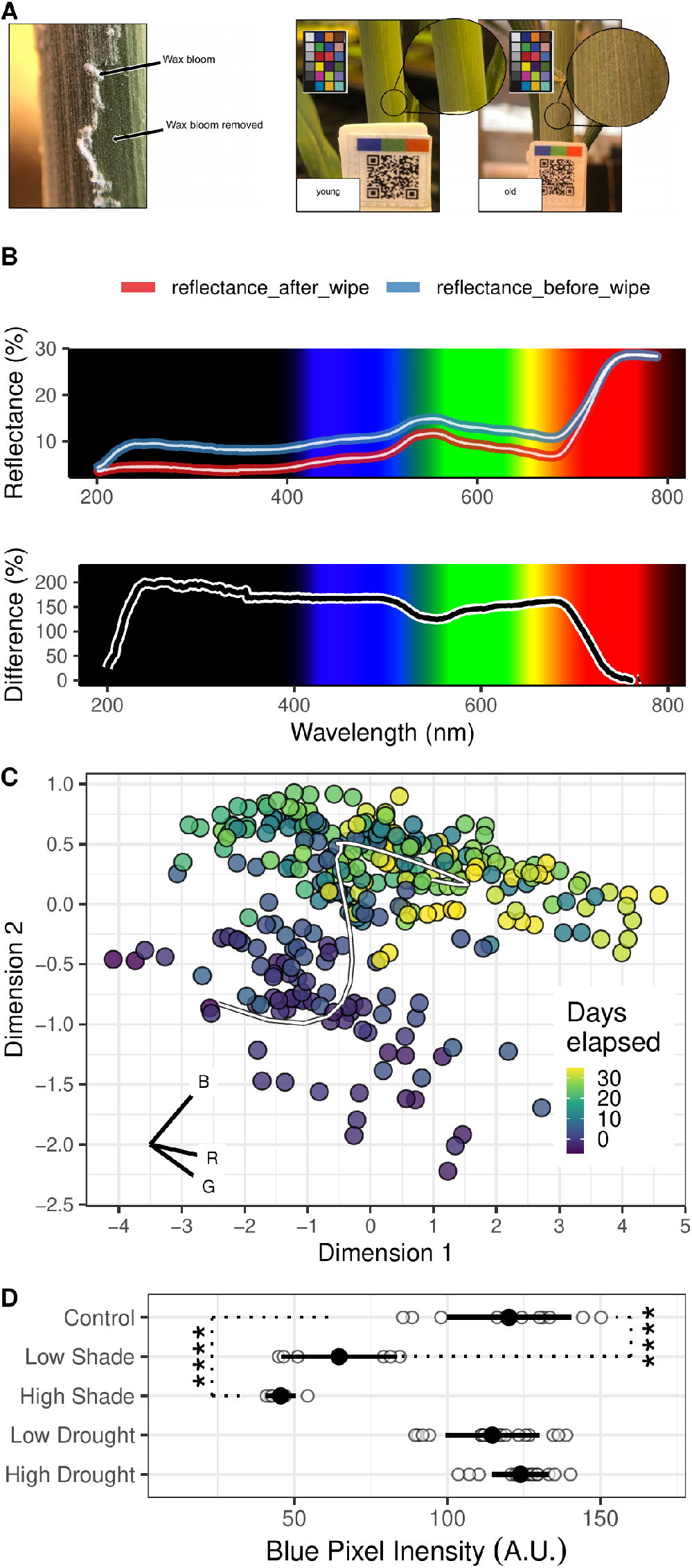
Visual phenotyping for sorghum wax A. Left: visual comparison of sorghum tissue with intact wax bloom and with wax bloom removed, right: visual comparison of wax covered and wax-less sorghum tissue alongside the color checker apparatus used for normalizing photographic measurements. **B** Line graph showing the reflectance (top) and difference in reflectance (bottom)between wax-covered and wax-less tissue from 200-800nm. **C** Scatterplot representing principal components analysis corresponding to pixel intensity data collected prior to and throughout wax extrusion on growing sorghum plants. The trend line represents the mean dim1 dim2 values over time. The inset ordination plot represents the relative contribution of each color channel to the variance observed in the data set. **D** Scatterplot showing blue color channel pixel intensity, which reflects wax accumulation, for photos of sorghum leaf sheath tissue (Tukey test, **** p-value ≤ 0.0001). White points represent individual pixel intensity values, and black points represent average pixel intensity for each treatment. Higher pixel intensity values indicate greater wax coverage. Error bars represent standard deviation of n = 12, n = 18, and n = 6 replicates for control, drought, and shade treatment groups, respectively.

## 4. Discussion

The primary aim of this study was to expand our understanding of sorghum leaf sheath wax extrusion and the mechanisms underlying the its response to different environmental stimuli. We used chemical and photographic analyses to compare wax accumulation on sorghum plants subjected to drought and simulated shade stress as well as RNA sequencing and differential expression analyses to compare gene expression between samples not extruding wax and samples in which wax extrusion had been induced. Here, we integrate these data with literature to discuss differences in sorghum leaf sheath wax load and composition in response to environmental stimuli and between sorghum varieties (section 4A), as well as a potential signaling pathway for wax bloom induction (section 4B), and the potential of smartphone-based phenotyping as a tool to further examine that hypothetical pathway (section 4C).

### A. Sorghum leaf sheath wax coverage and composition is altered under high intensity drought or simulated shade stress and varies by cultivar

We used GC-MS (Fig. 1A, Fig. 2A and B) to explore the dynamics of wax bloom extrusion over two timescales. On a long timescale, our greenhouse experiments highlighted modulations in wax bloom extrusion that occurred over approximately four months. GC-MS measurements from the greenhouse experiments discussed above showed that sorghum plants grown under high intensity drought produced significantly more wax (64 *μg cm*^−2^) than control plants (38 *μg cm*^−2^), and plants grown under high intensity simulated shade produced significantly less wax (17 *μg cm*^−2^) than control plants after approximately four months of growth. These findings align with previously reported wax coverages for irrigated and non-irrigated sorghum from ~38-215 *μg cm*^−2^, as well as previous work demonstrating that wax accumulation is influenced by light and drought in plant species ranging from agricultural crops to the bryophyte species *Phsycomitrella patens* (29, 31, 32, 38–44, 57–66). It should be noted that the wax coverages reported in this work were on surfaces of the cultivar BTX623, while the wax coverages reported in literature were on a variety of other cultivars. No literature data was available for wax coverage on shade-grown sorghum. The variation beteween the wax loads reported in this work and those reported in the literature may be accounted for by the diversity in wax load and composition that can occur on sorghum grown under different environmental conditions or between cultivars and underscore the dynamic nature of this trait with regard to likely both genotype and environment on short (72 hour) and long (120 day) timescales. Together, these results illustrate the long-timescale wax extrusion dynamics that occur on the order of months.

Our light-based induction experiments highlighted modulations in wax bloom extrusion that occurred over the course of 72 hours. GC-MS measurements from light-based induction experiments showed that there was minimal wax coverage (4 *μg cm*^−2^) on sorghum leaf sheath tissue harvested from within the culm of a mature sorghum plant, but significant wax accumulation (14 ± 6 *μg cm*^−2^) occured after 72 hours of light exposure (Fig. 2A and B). Since epicuticular waxes in plants primarily serve functions related to protection against drought and photodamage (16, 55), a potential explanation for the near absence of epicuticular waxes on sorghum tissue covered by overlapping leaf sheath layers is that these layers protect the developing sorghum tissue within the culm from desiccation and photodamage, reducing the immediate need for wax coverage. It may be advantageous to delay the extrusion of metabolically expensive compounds like the VLCFAs found in sorghum wax until their functional necessity peaks. In contrast, leaf sheath tissue that has emerged from the culm is no longer protected from desiccation and photodamage by the overlying tissue and thus benefits from the presence of wax blooms. The contrasting stressors faced by pre-emergent (metabolic limitations) and post-emergent tissues (UV light and drought) align with the timing of wax emergence, which appears to coincide with its protective role against drought and photodamage. Together these findings further illustrate the dynamic nature of wax extrusion, though on the timescale of days, rather than months. Together, these results illustrate the short-timescale wax extrusion dynamics that occur on the timescale of days.

Increased wax load under drought conditions has been documented not only in sorghum but also in various other species including tobacco, maize, alfalfa, wheat, and brassica species (25, 29, 35, 37, 62, 67). For example, Lee et al. reported that leaf wax increased under drought conditions in broccoli, correlating with enhanced drought tolerance. Similarly, Ni et al. reported increased wax content in certain alfalfa genotypes under drought stress, which was linked to improved drought resistance. Islam et al. found that the accumulation of leaf wax in rice was associated with drought tolerance, and Meeks et al. found that genetic variation in in maize epicuticular wax was associated with differences in drought tolerance. Overall, these data highlight the widespread role of dynamic wax production as an adaptive mechanism for enhancing drought tolerance across a variety of plant species.

Several of the genes in our expression dataset, such as HY5, SAPK9, CBL-interacting protein kinase 2, DEHYDRATION INDUCED-19, LTP3, and HVA22, are also associated with drought tolerance (5, 6, 68–73). Given that wax blooms are associated with protection against environmental stressors like sunlight and drought, and the differentially expressed genes mentioned above are associated with tolerance to these stressors, it is plausible that these genes regulate expression of wax-related genes. In turn, these pathways may confer drought tolerance via wax extrusion. While there is disagreement about whether or not epicuticular waxes form a transpiration barrier capable of significantly impacting drought tolerance (74, 75), previous studies have demonstrated that wax blooms can reduce non-stomatal water loss and are associated with drought tolerance in several species (76–80). Beyond the well-documented role of epicuticular waxes in drought tolerance, it is perhaps less well-known that the dynamic induction and modulation of wax extrusion appear to be a widespread strategy for managing envrionmental stressors. (31, 66, 67, 81, 82).

Our GC-MS results show that the primary chemical constitutents of grain sorghum (var. BTX623) leaf sheath wax blooms are C30, together with significantly smaller amounts of C28 fatty acids (Fig. 1B). These findings agree with previous work demonstrating that the wax blooms of sorghum (var. BTX623) are comprised of approximately 95% free fatty acids, but contrast with the previous findings that abundance of C28 fatty acids is greater than that of C30 (66, 83). It should be noted that the wax chemistries reported by Avato et al. correspond to waxes on leaf tissue rather than leaf sheath tissue which was used for our analyses. Our findings also contrast with the findings of Chemelewski et al., 2023 which demonstrated that C28/C30 fatty alcohols are the primary chemical constituents of wax blooms occurring on the leaf sheaths of growing bioenergy sorghum plants (vars. Wray, R07020, and TX08001) (65). Together, these findings suggest that different sorghum cultivars (i.e., bioenergy versus grain sorghum) and that sorghum tissue types exhibit distinct wax chemistry, as has been observed in other species (64, 65, 84).

Our GC-MS results from the greenhouse experiments (Fig. 1B) show significant differences in wax composition between plants grown under high intensity drought and shade conditions compared to control conditions. Under high intensity drought conditions, plants showed significantly lower abundance of C30 fatty acids, and significantly higher abundance of C28 fatty acids compared to control plants. High intensity shade treatments led to significant increases in the relative abundance of both C28 and C30 alcohols. These data align with past findings that surface wax composition and fatty acid abundance are modulated under stressful conditions in other species (85–88). However, the specific alterations in VLCFA and VLCOH chain length that we observed have not previously been reported. Our data suggest that drought conditions increase the relative abundance of shorter (C28) chain VLCFAs compared to longer (C30) chain VLCFAs while shade conditions increase the relative abundance of C28 and C30 alcohols. The functional purpose of these alterations in VLCFA and VLCOH remains unclear, as do the specific functions of many cuticular wax components, but the dynamic nature of many species’ wax mixtures in response to environmental stimuli and direct experimental evidence (62, 85–89) strongly suggest specific, if complex, roles for different wax chemicals in tuning the function(s) of cuticular waxes as a whole.

### B. A draft signaling pathway for wax bloom induction in sorghum based on differentially expressed genes during wax extrusion

The light responsivity of wax bloom extrusion has been demonstrated by previous experiments showing that light exposure can induce wax blooms on tissue excised from beneath three to four layers of overlying tissue (Fig. 2B) (54). Intriguingly, minimal wax induction is observed when such tissue is maintained in darkness, highlighting the importance of light as a stimulus. As with virtually any physiological process, possessing the proper molecular machinery is a pre-requisite for responding to a stimulus, which, in the case of sorghum wax bloom extrusion, must consist of at least (i) light-responsive signaling mechanisms, (ii) wax biosynthesis, and (iii) wax transport. Our RNA-seq data (Fig. 2C) suggest that during light induction of wax blooms, light-responsive gene expression generates the molecular machinery (transcription factors, enzymes, transporters) responsible for the production and extrusion of epicuticular waxes in response to light. Below, we discuss four features of a hypothetical signaling pathway for wax bloom induction.

We found several key light perception and signaling genes upregulated in our dataset, including the photoreceptors phyA, PHOT2, and UVR8, which are known to play roles in light perception (90–98). Additionally, light signaling genes such as ELONGATED HYPOCOTYL (HY5), PHYTOCHROME INTERACTING FACTORS (PIFs), and transcription factors (TFs) similar to several MYB domain proteins were also implicated by our gene expression data. HY5 is a convergence point for several signaling pathways in plants (99–103) and is involved in coordination of the photoprotective response in response to light fluctuation (104). HY5 also mediates the signal transduction of several photoreceptors, including some that were identified in our gene expression dataset (phyA, UVR8), and interacts with and regulates a multitude of transcription factors and signaling proteins including some from our dataset (105–108). Based on these observations, it appears that HY5 is likely a key, central component of the signaling cascade mediating the induction of wax blooms in sorghum. In line with these data, we hypothesize that the signaling pathway mediating the induciblity of sorghum wax blooms is initiated by photoreceptors that interact with TFs like HY5. In turn, HY5 may interact with other TFs and/or promoters of wax transport genes like ABCG11 and LTPs (upregulated in our dataset), enhancing their expression during wax extrusion.

During our literature searches, we identified a variety signaling interactions between genes in our wax induction dataset. For example, phyA has been shown to directly influence UNE10 (PIF8), as well as IAA17 (107, 109, 110). Additionally, phyA negatively regulates PIF5, which in turn influences IAA19/29, a protein that has been shown to interact with IAA17 to inhibit cell elongation during the shade avoidance response (105, 111, 112). Chemelewski et al. found that the onset of wax extrusion was correlated with the termination of cell elongation on sorghum internodes (65). Given that IAA17 has been implicated as a regulator of cell elongation in response to different light conditions (113, 114), we speculate that the phyA/PIF5/IAA19/29/IAA17 signaling module may be involved in regulating the onset of wax extrusion.

We also obsered differential expression of regulators of MYB activity including the RING-type E3 ligase MYB30-INTERACTING E3 LIGASE 1 (MIEL1) and TF NIGHT LIGHT-INDUCIBLE AND CLOCK-REGULATED1 (LNK1) during wax induction (115, 116). Notably, MIEL1 has been shown to negatively regulate wax biosynthesis in Arabidopsis, possibly through its interaction with several wax-related MYB TFs including MYB30 and MYB96 (117). Interestingly, this suggests that the time point at which we sampled, 72 hours after exposure to light, may have occured during or after the arrest of wax biosynthesis in which MIEL1 has turned off TFs similar to MYB96 and MYB30. Others have looked at snapshots of gene expression during different periods of wax induction. In a recent paper by Chemelewski et al., gene expression data collected during sorghum internode development was used to identify candidates involded in regulating sorghum leaf sheath wax extrusion (65). In contrast to the candidates in our expression dataset which were primarily involved in light/hormone signaling and wax transport, many of the candidates identified by Chemelewski et al. have been associated with wax biosynthesis (KCS, KCR, ECR, MAH, LACS) or regulation of wax biosynthesis (MYB94/96/30/60) rather than transport (118–129). Though these differences could be in part or whole to genotypic differences, it seems highly likely that there is a temporal aspect on the scale of days to the induction process. Despite the differences in gene expression discussed above, the ATP-binding cassette transporter ABCG11 was associated with wax extrusion in both our and Chemelewski et al., expression datasets. The involvement of ABCG11 in wax transport has been documented extensively (130–135). However, additional studies focused on gene expression from the onset of stem internode cell elongation through the termination of wax extrusion across multiple sorghum cultivars will be necessary to clarify the full regulatory network underlying wax extrusion in sorghum.

There remain gaps in our signaling model that will require elucidation before the regulatory network underlying wax bloom induction is fully understood. Given that HY5 is involved in the integration of both light and hormone signaling (108), and that certain hormones have been shown to influence epicuticular/cuticular waxes (136, 137), we hypothesize that hormone signaling pathways may account for the missing components of the regulatory network. Within our gene expression dataset there were a variety of differentially expressed hormone-related genes. Auxin-inducible genes like INDOLE ACETIC ACID 14 and 17 (IAA14 and IAA17), and 5NG4 (138, 139) were differentially expressed during wax induction. Abscisic acid (ABA) signaling genes, including CBL-INTERACTING PROTEIN KINASE 2, abscisic stress ripening protein-1-like, GTE9, and others (6, 72, 140, 141) were also differentially expressed during wax induction. In addition to genes whose expression is influenced by a single hormone, there were also genes in our gene expression dataset (SAPK9 and bZIP11) whose expression is influenced by multiple hormones, specifically ABA and auxin, as well as genes which have been implicated in both hormone and light signaling pathways (SAPK9) (142, 143). Interestingly, previous work has demonstrated that HY5 promotes the expression of IAA14 and other auxin-related genes. The presence of genes related to both hormone and light in our differential expression dataset indicate that there may be crosstalk occuring between multiple signaling pathways during wax induction. Furthermore, previous work has shown that the class sucrose nonfermenting protein kinase 2 subclass III (SNRK2.3), stress/ABA-activated protein kinase 9/10 (SAPK9/10) interact with various basic leucine zipper (bZIP) proteins related to bZIP11, an auxin/ABA-inducible TF that was differentially expressed during wax induction (144). The SNRK2.3 protein SAPK9 may phosphorylate bZIP11 and CBL-interacting protein kinase, as the closely related SNRK2.3 SAPK10 has been shown to interact with CBL-interacting protein kinases and bZIP TFs (71, 144, 145). The molecular factors responsible for connecting the signaling modules discussed above (both light signaling and hormone signaling, etc) to wax-related machinery are yet unclear and seem complex. However, the signaling components outlined earlier in this section (HY5, phyA/PIF5/IAA19/29/IAA17, and termination by MIEL1) serve as starting points from which future researchers can expand our understanding of the signaling network underlying the induction of wax bloom extrusion.

### C. Wax bloom extrusion dynamics can be monitored using smartphone-based phenotyping

At the end of the greenhouse experiment, wax coverage on the shade treatment group was visibly different compared to the control group, but the drought treatment was not. Results of both the GC-MS (Fig. 1A) and photographic(Fig. 3B) analyses showed that control and drought treatment groups exhibited significantly greater wax accumulation compared to shade treatment groups. Although we were unable to detect the significant differences in wax coverage between control and drought treatment groups that were detected via the GC-MS method, significant differenes in wax coverage were detected between both shade treatment groups and the control group using the photographic method. One potential explanation for our inability to detect differences between the drought treatment and control groups is that the dynamic range of color-based detection methods has a lower limit of linearity than that of GC-MS. While this photographic method may have fewer capabilities than more cost and labor intensive analytical methods like GC-MS, it seems effective only at detecting the visual differences between sparse and moderately heavy wax blooms, as opposed to the difference between moderately heavy and very heavy wax blooms. Even so, this drawback may not be entirely problematic since the mutation of wax-related genes often results in bloomless or sparse bloom phenotypes (25, 146–149), and detecting differences between wildtype (control) wax load and sparse (bloomless/sparse bloom mutant) is within the capabilities of a smartphone being used as a high throughput screening apparatus for assaying wax-related genes.

Our photographic data collection and analysis pipeline begins with a researcher capturing close-up photographs of sorghum leaf sheaths with a color normalization device (calibrite color checker) in frame. Next, the photographs are uploaded into our analysis application where the RGB values of each photograph are adjusted such that the pixels associated with each reference swatch of the color normalization device are the identical. In doing this, it is possible to correct for differences in lighting between different photographs. Once normalized, the experimentalist selects the region of interest on the photograph using a drag and drop box within the analysis application. For the pixels within the selected region, the application determines which intensity value occurs most frequently for the blue color channel (i.e., the mode) and records that value. Our choice to use intensity values corresponding to the blue color channel of our photographic data was based on the data presented in section 3C (Fig. 3A-D), where our results showed that blue light is reflected more strongly than other wavelengths within the visible region of the spectrum and that the variance in our preliminary photographic dataset was primarily attributed to the blue color channel during wax extrusion. Our photograph analysis application also converts photographic measurements into 0-255 values allowing for semiquantitative measurement. The semiquantitative nature of this system may enable detection of phenotypes that are more subtle than those that could be detected with the naked eye and may also enhance uniformity in measurements between researchers. Using this system, a pair of researchers could make daily measurements and analyze photographic data from approximately three to four dozen lines over the course of a three month field trial. Thus, our photographic data collection and analysis system provides a rapid and semiquantitative method for collecting wax bloom phenotypic data that would enable a group of researchers to test many, or all, of the candidate genes discussed above.

## 5. Conclusions

This study provides insights into the dynamics of epicuticular wax bloom extrusion in sorghum and highlights a smartphone-based phenotyping system as a potential tool for investigating candidate genes identified. By comparing wax load on sorghum grown under different environmental conditions, we demonstrated that wax bloom production is altered in response to light and drought stress. Using an *in vitro* wax induction protocol we showed that light is a necessary stimulus for significant wax extrusion. Using gene expression data from wax induction experiments we identified candidate genes potentially involved in regulating wax extrusion, showed that wax extrusion appears to be coordinated with termination of cell elongation, and highlighted that MIEL1 may be involved in terminating wax biosynthesis during wax extrusion. Additional studies will be needed to clarify the full regulatory network underlying wax bloom induction. Our gene expression data provide a list of molecular candidates associated with wax bloom extrusion, and our smartphone-based phenotyping system provides a means through which future researchers can study them. Finally, we considered the sorghum genes differentially expressed in response to light stimulus and examined their orthologs in maize. A wide variety of sorghum genes, including thaumatin-like and BURP- and MYB-domain containing proteins, had orthologs in maize that were not upregulated in response to light, suggesting that comparative genomics may be an important component in future studies aimed at engineering wax blooms in maize. This work has highlighted the complex interplay between environmental factors and epicuticular wax production in sorghum and introduced a tool that can help us futher deepen our understanding of a trait that shows potential in the engineering of more resiliant crops.

## 6. Acknowledgements

The authors wish to acknowledge the support of Zhanguo Xin, Al Oyler. We also wish to acknowledge the University of Minnesota Duluth Chemistry and Biochemistry Department, UMD Undergraduate Research Opportunities Program, UMD Summer Undergraduate Research Program, UMD Chemistry Master’s program, UMD Moses Passer Graduate Fellowship, UMD Siders Graduate Fellowship for providing the funding and facilities to make this research possible. We collectively acknowledge that the University of Minnesota Duluth is located on the traditional, ancestral, and contemporary lands of Indigenous people. The University resides on land that was cared for and called home by the Ojibwe people, before them the Dakota and Northern Cheyenne people, and other Native peoples from time immemorial. Ceded by the Ojibwe in an 1854 treaty, this land holds great historical, spiritual, and personal significance for its original stewards, the Native nations, and peoples of this region. We recognize and continually support and advocate for the sovereignty of the Native nations in this territory and beyond. By offering this land acknowledgment, we affirm tribal sovereignty and will work to hold the University of Minnesota Duluth accountable to American Indian peoples and nations.

## Notes

https://github.com/thebustalab/wax_bloom_dynamics

### Competing Interest Statement

The authors have declared no competing interest.

